# BMP9-mediated regulation of endothelin-1 requires integrated SMAD1/5 and SMAD2/3 signaling

**DOI:** 10.64898/2025.12.24.696445

**Authors:** Jana Bagarova, Shreya Sangam, Luca Troncone, Ying Zhong, Lillian C. Worst, Greg R. Gipson, Ivana Nikolic, Samuel Paskin-Flerlage, S. Paul Oh, Fernando Rodríguez-Pascual, Paul B. Yu

## Abstract

BMP9 is a pleiotropic growth factor cytokine with diverse roles in vascular development, homeostasis and disease. BMP9 regulates a broad array of vasoactive molecules that mediate endothelial and mural cell function, including ET-1, a potent vasoconstrictor, regulator of cell growth and fibrosis, and therapeutic target for pulmonary arterial hypertension (PAH). Consistent with its pleiotropic activities, BMP9 is unique in being able to recruit both BMP-responsive SMAD1/5/9 and TGFβ-responsive SMAD2/3 signaling effectors in endothelial cells, however, the physiologic significance of activating both pathways remains incompletely defined. We investigated the role of SMAD1/5/9 vs. SMAD2/3 signaling in BMP9-mediated regulation of ET-1, using primary and immortalized human and murine microvascular endothelial cells, with conditional knockout, small molecule inhibitor and siRNA strategies. BMP9-mediated expression of ET-1 requires coordinated activation of SMAD1/5/9 and SMAD2/3 effector pathways, both downstream of BMPR2, ALK1, and ENG. Analysis of the *ET-1* promoter revealed that BMP9 requires, in addition to a SMAD3/4 binding site sufficient for TGFβ1-mediated transcription, a novel putative SMAD1/5 binding motif. BMP9-mediated regulation of endothelial ET-1 requires coordinated activation of both SMAD1/5 and SMAD2/3 downstream of ALK1, integrated at the promoter level, representing a non-canonical signaling motif linking BMP9 to a critical effector of vascular tone and remodeling in PAH and related vascular syndromes.

**Translational Perspective:** BMP9 has recently emerged as a novel therapeutic target with an ongoing Phase 2 clinical trial testing an anti-BMP9 antibody for efficacy in PAH (NCT06137742). The mechanisms by which BMP9 contributes to pulmonary vascular disease are thought to include its regulation of Endothelin-1 (ET-1), an established therapeutic target in PAH. We show that BMP9 potently regulates ET-1 transcription via transactivation of TGFβ1 effector SMAD3 via BMP receptors BMPR2 and ALK1, suggesting that pleiotropic effects of this growth factor cytokine result from non-canonical activation of BMP and TGFβ signaling pathways, with implications for its roles in physiology, and the rationale for its therapeutic modulation for PAH.

## INTRODUCTION

Endothelin-1 (ET-1), a potent vasoconstrictive peptide,^1,2^ is expressed primarily in vascular endothelial cells (ECs), and to a lesser extent in vascular smooth muscle cells (SMCs), fibroblasts, and cardiomyocytes. Beyond its well-established vasopressor function, ET-1 regulates diverse aspects of vascular homeostasis, angiogenesis, extracellular matrix production, fibrosis, inflammation, and oxidative stress.^3^ These wide-ranging effects on vascular structure and physiology, and observations that ET-1 exerts potent mitogenic effects on vascular ECs, SMCs, and fibroblasts^4^ are all consistent with its critical role in pulmonary vascular remodeling and pulmonary arterial hypertension (PAH). Vascular endothelium-derived ET-1 appears to coordinate these processes, and is in turn responsive to a wide variety of stimuli including insulin, thrombin, angiotensin, vasopressin, hemodynamic shear stress, hypoxia, nitric oxide, and transforming growth factor-β (TGFβ) signaling.^2^ Circulating levels of ET-1 are elevated in patients with PAH, and predictive of invasive hemodynamics, exercise function, and mortality.^5–10^ ET-1 is highly expressed in the endothelium of vascular lesions in PAH lungs by immunostaining and in situ hybridization, suggesting important roles in vascular pathology.^11^ ET-1 receptor antagonists (ERA) are a cornerstone of first-line therapy in PAH, improving functional status, hemodynamics, and a combined endpoint of morbidity and mortality in a long-term clinical trial.^12^

Previous work has highlighted bone morphogenetic protein 9 (BMP9, encoded by *growth and differentiation factor 2*, *GDF2*) as being relevant to PAH and other vascular syndromes. Vascular ECs are highly enriched for the expression of proteins forming the BMP9 and BMP10 signaling complex, including the BMP type II receptor (*BMPR2*), BMP type I receptor ALK1 (*ACVRL1*), and BMP co-receptor Endoglin (*ENG*).^13–17^ Various loss-of-function mutations in *BMPR2*, *ACVRL1*, *ENG*, *GDF2, and BMP10*—encoding an endothelial BMP signaling complex—are implicated in heritable PAH (HPAH) and hereditary hemorrhagic telangiectasia (HHT) syndromes.^18–23^ BMP9 is a ligand that is produced in hepatic stellate cells of the liver and is present at physiologically active concentrations in the circulation,^18,24^ The abundance of BMP9 in the circulation, its effects in promoting endothelial homeostasis and regulating vascular phenotypes have suggested its role as a circulating vascular quiescence factor.^18,25^ Consistent with this notion, circulating levels of BMP9 are markedly diminished in portopulmonary hypertension and in hepatopulmonary syndrome,^23,26–28^ suggesting acquired deficiency of BMP9 may contribute to pulmonary vascular disease in severe liver disease. BMP9 potently modulates growth, survival, migration, and inflammation in vascular ECs.^14,18,29^ Classic BMPs achieve downstream functions by activating BMP-responsive SMAD effector proteins 1, 5, and 9 (SMAD1/5/9), whereas classic TGFβ ligands activate TGFβ-responsive SMADs 2 and 3 (SMAD2/3). BMP9 and BMP10 appear to activate both SMAD1/5/9 and SMAD2/3 in ECs,^14,30–32^ however, the mechanisms by which BMP9 recruits both BMP and TGFβ-mediated signaling, and the functional consequences of dual activation of BMP and TGFβ signaling have yet to be fully defined.

Prior studies demonstrated TGFβ regulates ET-1 in ECs by activation of canonical TGFβ effector SMAD3 by the TGFβ type 1 receptor ALK5.^33^ TGFβ-mediated expression of ET-1 was found to depend on the cooperative binding of SMAD3/4 to a SMAD binding element (SBE) along with an activator protein 1 (AP-1) site in the *ET-1* promoter.^33–35^ Subsequent reports described the regulation of ET-1 by BMP9,^29,36,37^ suggesting dysregulated BMP9 signaling associated with various vascular syndromes could affect vascular tone and/or remodeling via altered ET-1 expression. In fact, the protection of *Gdf2^-/-^* mice from experimental PH has been attributed, in part, to associated loss of BMP9-mediated ET-1 expression in these animals.^38,39^ Moreover, exogenous BMP9 potently induces vasoconstriction in chick chorioallantoic membranes, and in turn is inhibited by the endothelin receptor antagonist (ERA) bosentan, demonstrating an ET-1-dependent mechanism for BMP9-mediated vasoconstriction.^39^ Based on this work and subsequent work by our group demonstrating that BMP9 is essential for arterialization of ECs associated with vascular remodeling in experimental PH, and a potential mechanism of action for sotatercept,^40^ selective inhibition of BMP9 is now being tested in a Phase 2 clinical trial for the treatment of PAH (NCT06137742).^41^

In the present study, we sought to determine the signaling requirements for BMP9-mediated regulation of ET-1 and found activation of SMAD2 and SMAD3 by BMPR2 and ALK1 are essential for expression of *EDN1* (encoding ET-1). In analyzing the *EDN1* promoter, we found that a previously identified TGFβ-responsive *cis*-acting SMAD2/3 binding element^42^ as well as a novel SMAD1/5 binding element were both required for efficient BMP9-mediated *EDN1* transcription. Regulation of *EDN1* by BMP9 appears to require the cooperation of canonical BMPR2/ALK1-SMAD1 and non-canonical BMPR2/ALK1-SMAD3 signaling, potentially representing a novel motif for regulation of other vasoactive molecules in endothelium The properties of BMP9 as multi-functional BMP/TGFβ agonist may contribute to its pleiotropic and context-sensitive functions as both a quiescence factor and driver of vascular pathology.

## METHODS

### Cell culture

Bovine aortic endothelial cells (BAECs) obtained from Lonza (Walkersville, MD) were cultured in Dulbecco modified Eagle’s medium (DMEM) with high glucose supplemented with 10% fetal calf serum, glutamine and antibiotics. Human pulmonary artery endothelial cells (HPAECs) were cultured in EGM-2 complete growth medium (Lonza). Telomerase immortalized microvascular endothelial (TIME) cells obtained from ATCC (Manassas, VA) were cultured in vascular cell basal medium (VCBM) supplemented with vascular endothelial growth factor (VEGF). Mouse lung microvascular endothelial cells (MLECs) were isolated as previously described, from mice harboring two conditional knockout *Bmpr2* (*Bmpr2^flox/flox^*)^43^ or *Acvrl1* (*Acvrl1^flox/flox^*) alleles.^13^ MLECs were purified using rat anti-mouse CD31 antibody (Clone MEC13.3, BD Pharmingen), and superparamagnetic beads conjugated to sheep anti-rat IgG Fc (Dynabeads, Life Technologies), and cultured in DMEM supplemented with 20% fetal calf serum, glutamine, penicillin, streptomycin, non-essential amino acids, 100 µg/mL heparin (Sigma, St. Louis, MO), and EC growth supplement (100 µg/mL, BD Biosciences).

### Ex-vivo recombination and generation of *Bmpr2* and *Alk1* homozygous knockout endothelial cells

The *ex-vivo* excision of homozygous floxed murine *Bmpr2* (*Bmpr2*^flox/flox^) or *Acvrl1* (*Acvrl1*^flox/flox^) sequences in conditional knockout MLECs was achieved by culturing with Adenovirus expressing Cre recombinase (Ad.Cre, Vector Labs) at a multiplicity of infection (MOI) of 20, whereas control cells were treated with equal MOI of Adenovirus expressing GFP (Ad.GFP). Disruption of *Bmpr2* or *Acvrl1* in ECs was confirmed by quantitative real-time PCR and/or immunoblot (monoclonal anti-BMPR2, BD Biosciences) as previously described.^13,44^ In the case of *Acvrl1^-/-^*, MLECs were isolated to clonal purity by limiting dilution.^13^

### Promoter Constructs

Luciferase reporter constructs driven by human *ET-1* promoter sequence varying in length from 0 to -1.5 kbp were generated as previously described.^33–35^ Point mutations in AP-1, SBE1 and SBE2 sites,^34^ as well as a novel construct eliminating the NF-1 binding site were introduced into the -650 bp segment of the wild-type human *ET-1* promoter using the QuickChange Site Directed Mutagenesis kit (Agilent Technologies, Santa Clara, CA). CAGA-Luc, a TGFβ signaling luciferase reporter (CAGA-luc) consisting of 12 tandem repeats of the upstream Smad3-binding element from human *PAI-1* promoter linked to a viral minimal promoter and firefly luciferase,^45^ and BRE-Luc, a BMP responsive signaling luciferase reporter (BRE-luc) construct consisting of two tandem repeats of the Smad elements from murine Id-1 linked to a minimal promoter and luciferase were kindly provided by Peter ten Dijke (Leiden University Medical Center).^46^

### Luciferase reporter assay

Cultured ECs were transfected in 96-well plates with 0.1 μg of plasmid encoding firefly luciferase driven by various *ET-1* promoter segments, and 0.1 μg control plasmid encoding Renilla luciferase driven by the CMV promoter. Plasmid DNA was delivered using PolyJet transfection reagent (SignaGen Laboratories, Rockville, MD) overnight, followed by 8 h serum deprivation, and subsequent stimulation with recombinant human BMP9 or TGFβ1 (R&D Systems, Minneapolis, MN), respectively, and analyzed using the Dual Luciferase Kit (Promega) with a SpectraMax L luminometer (Molecular Devices, Sunnyvale, CA). Relative *ET-1* promoter activity was calculated as the firefly luciferase divided by Renilla luciferase activity.

### siRNA transfection

Small interfering RNA (siRNA) were used to inhibit expression of BMP and TGFβ type I and II receptors, and ENG. Validated siRNA duplexes specific for human *BMPR2*, *ACVR2A*, *ACVRL1*, *ENG*, and *ALK5* mRNA were obtained from Applied Biosystems (Supplemental Table 1) and transfected (5-30 nM) into cultured human pulmonary arterial endothelial cells (HPAECs) using PepMute transfection reagent (SignaGen) according to the manufacturer’s protocol. BAEC, HPAEC, or TIME cells were transfected with siRNA for 24 h, deprived of serum for 16 h, and stimulated with BMP9 or TGFβ1 for 3 h.

### Quantitative RT-PCR and ELISA

Total RNA was isolated using Isol RNA isolation reagent (5-PRIME) according to the manufacturer’s protocol. One μg of total RNA was subjected to reverse-transcriptase reaction for 1 h at 37°C using SuperScript II reverse transcriptase (Invitrogen). Relative cDNA levels were assayed by quantitative real time PCR (Kapa SYBR FAST Universal qPCR Kit, Kapa Biosystems). Secreted ET-1 peptide levels were determined using a sensitive ELISA (Big Endothelin-1 assay ADI-900-022, Enzo Life Sciences) according to the manufacturer’s protocol.

### Plasma samples

Plasma samples were taken from rats developing experimental PH, and control rats according to the following protocol. Adult male Sprague Dawley rats (6-8 weeks old, 150 to 170 g, Charles River Laboratories) were exposed to SU5416 (20 mg/kg x 1 dose s.c.) and 3 weeks of normobaric hypoxia (FIO2=0.10), followed by 3 weeks of normoxia, as well as age-matched normoxic (Nx) control rats, were used to obtain plasma for measurements of plasma ET-1 levels. Rats were anesthetized with isoflurane (2.5% induction, 0.5 to 1.5% maintenance), and right ventricular (RV) pressures were measured by a minimally invasive closed chest approach using a curved tip 2 French pressure transducer catheter (Millar, SPR-513) inserted into the RV through the right internal jugular vein. At completion of study, animals were euthanized with sodium thiopental (200 mg/kg i.p. x 1) and blood sampled via cardiac puncture and centrifuged (1500 g x 30 min with EDTA). Animal studies were performed under an approved protocol with oversight from the Massachusetts General Hospital Institutional Animal Care and Use Committees. Animals were housed at 24°C in a 12-hour light/12-hour dark cycle where food and water were accessible ad libitum. Plasma levels of ET-1 were measured also using ELISA kit ADI-900-022 (Enzo).

### BMP and TGFβ signaling inhibitors

BMP type I receptor (ALK1/2/3/6) kinase inhibitor, LDN-193189, was synthesized as previously described.^47,48^ Activin/TGFβ type I receptor (ALK4/5/7) kinase inhibitor, SB-431542, was obtained from Sigma.^49^ SIS3, a selective inhibitor of SMAD3 activation and function,^50^ was obtained from Tocris Bioscience. Recombinant ALK1 extracellular domain expressed as a soluble fusion protein with IgG Fc domain (ALK1-Fc), a neutralizing anti-BMP9 monoclonal antibody (MAB3209), and recombinant BMP inhibitor, Noggin, were obtained from R&D Systems. Following serum deprivation for 16 h, cells were incubated with specified inhibitor for 30 min at 37°C prior to ligand stimulation.

### Statistical analysis

Data are represented as mean ± SEM, unless otherwise noted. Statistical significance was determined using the Student’s unpaired *t* test or one-way ANOVA with Dunnet’s or Kruskal-Wallis tests for multiple comparisons. All analyses were performed using Prism 10.0 (GraphPad Software). An adjusted *p*-value of <0.05 was considered to be statistically significant. The scRNA-seq dataset (GSE169471, droplet-based data) was re-analyzed using the Seurat R package (4.0.5) for our study.^51^

## RESULTS

### *EDN1* is overexpressed in ECs of lungs with PAH

In UMAP analysis of single-cell RNA sequencing data from Saygin and colleagues,^51^ *EDN1* was expressed primarily in the EC clusters, and more abundantly expressed in explanted lungs from patients with PAH vs. control donor lungs (**Fig. 1A**). Within endothelial lineages, *EDN1* was identified as one of the most significantly upregulated genes in lungs from PAH, as shown by volcano plot (**Fig. 1B**). Consistent with its known role as a circulating biomarker of human PAH, ET-1 was elevated ∼5 fold in the plasma of rats undergoing 3 weeks of treatment with SU5416 and hypoxia (FIO2=0.10) followed by 3 weeks of normoxia, versus control Nx rats (**Fig. 1C**).

**Figure 1.**
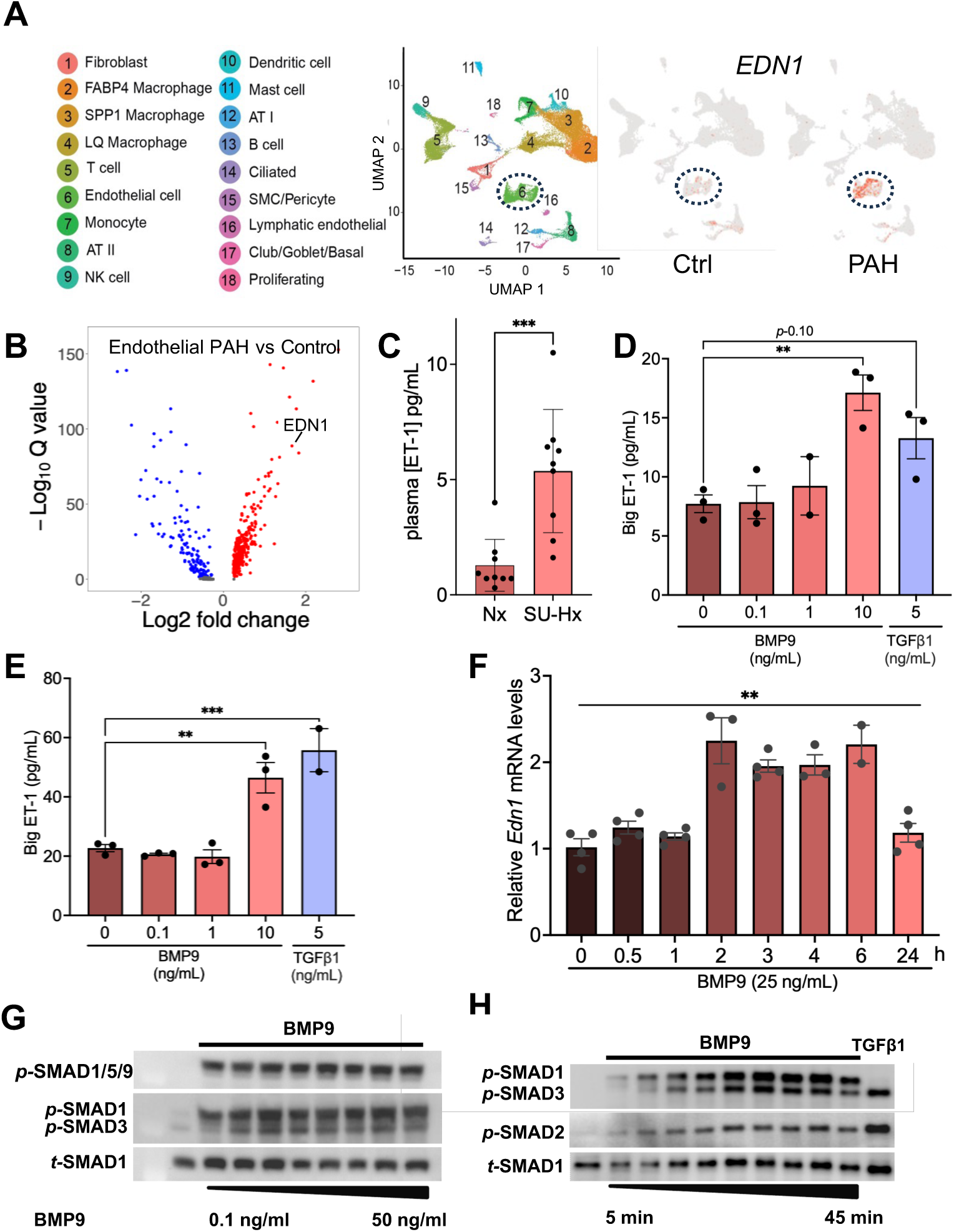
BMP9 induces expression of ET-1 peptide and *Edn1*, while activating SMAD1/5/9 as well as SMAD2/3. (A) UMAP plots depict distinct cell clusters in control (Ctrl) versus PAH explanted lungs identified by single cell RNAseq. *EDN1* expression is highly enriched in the endothelial cluster of cells in both control and PAH lungs, and is increased in the lungs of PAH versus control lungs (GSE169471). (B) Volcano plot depicts upregulated and downregulated genes in explanted lungs from patients with PAH versus non-diseased controls. *EDN1* is one of the most highly and significantly upregulated genes (GSE169471). (C) ET-1 levels were measured in plasma from normoxic (Nx) adult rats, and rats injected with SU5416 (40 mg/kg s.c.) and exposed to 3 weeks of hypoxia (FIO2=0.1) followed by 3 weeks of normoxia (SU-Hx). (D-E) Bovine aortic ECs (BAECs) were deprived of serum for 16 h and treated with varying concentrations of BMP9 (0.1, 1, 10 ng/mL) or TGFβ1 (5 ng/mL), and supernatants analyzed at (D) 6 h or (E) 24 h for presence of ET-1 peptide by ELISA, revealing dose-dependent effects. (F) Analysis of *Edn1* mRNA by qRT-PCR in BAECs stimulated with BMP9 (25 ng/mL) revealed time-dependent effects. (G) Immunoblotting of BAECs stimulated with varying concentrations (0-50 ng/mL) of BMP9 for 25 min demonstrated activation of SMADs 1/5/9 in a dose-dependent manner. (H) Immunoblotting of BAECs stimulated with BMP9 or TGFβ1 revealed kinetics of SMAD2 and SMAD3 activation. Values are mean ± SEM, n=5, \**p* ≤0.05 and \*\**p*<0.01 as compared to basal transcriptional levels, 1-Way ANOVA, Kruskal-Wallis Test.

### BMP9 induces ET-1 expression in vascular ECs and promotes the phosphorylation of BMP- and TGFβ-associated SMADs

Given previous work highlighting the role of ET-1 downstream of BMP9 and TGFβ1 signaling,^29,33–37^ we examined the role of BMP and TGFβ signaling in the regulation of ET-1 in endothelium. Stimulation of quiescent BAECs with either BMP9 or TGFβ1 led to accumulation of the full-length precursor of ET-1 (big ET-1) in culture supernatants detected at 6 and 24 h with a sensitive ELISA (**Fig. 1D-E**), with exposure to 10 ng/mL (400 pM) BMP9 yielding similar effects to 5 ng/mL (200 pM) TGFβ1. To assess BMP9-mediated transcriptional regulation of ET-1, quiescent BAECs were stimulated with BMP9, and *Edn1* mRNA levels measured by RT-PCR at varying time points (0.5 - 24 h), revealing sustained increases in *Edn1* mRNA from 2-6 h compared with controls (**Fig. 1F**). To identify potential signaling effectors mediating BMP9-induced expression of ET-1, phosphorylated SMAD 1/5/9, phosphorylated SMAD1 and SMAD3, phosphorylated SMAD2, and total SMAD1 were assessed by immunoblot in BAECs treated with varying concentrations of BMP9 or for varying times (**Figs. 1G-H**). BMP9 elicited phosphorylation of BMP-associated SMAD1/5/9 at lower concentrations than for SMAD3 (≥ 0.1 ng/mL or 4 pM vs. ≥ 0.25 ng/mL or 10 pM, **Figs. 1, S1A**) and activated SMAD1 with slightly faster kinetics than SMADs 2 or 3 (≥ 5 min. vs. ≥10 min., **Figs. 1H**, **Fig. S1B**). TGFβ1 (5 ng/mL or 200 pM, 25 min.) demonstrated potent activation of SMADs 2 and 3, without activating SMAD1/5/9. Similar observations were made in TIME cells, with BMP9 eliciting potent activation of SMAD1 and SMAD3, and TGFβ primarily inducing activation of SMAD3 (**Fig. S2A**).

### BMP9-mediated ET-1 expression involves BMP and TGFβ type I receptor activation

BMP9 is thought to recruit signaling in ECs via signaling complexes containing its high affinity BMP type I receptor ALK1, in preference to other BMP/TGFβ type I receptors ALK2-7.^52^ While ALK1 is thought to balance the effects of ALK5 in regulating angiogenic signaling functions of TGFβ, ALK1 can also function cooperatively with ALK5, being recruited in signaling complexes to achieve “lateral” signaling via both SMAD1/5/9 and SMAD2/3.^53,54^ We probed the contribution of type I BMP (ALK1, 2, 3 and 6) and TGFβ receptor (ALK4, 5, and 7) kinases in BMP9-mediated SMAD1/5/9 activation, SMAD2/3 activation, and ET-1 expression using small molecule inhibitors of BMP (LDN-193189) and TGFβ (SB-431542) type I receptors, respectively. A concentration of ≥ 500 nM LDN-193189 was required to inhibit BMP9-mediated phosphorylation of SMAD1, whereas a concentration of ≥ 125 nM was sufficient to abrogate SMAD3 activation (**Figs. 2A, S2B**) in BAECs. Similarly, higher concentrations of LDN-193189 (≥ 500 nM) were required to block activation of SMAD1 while 125 nM was sufficient to block the activation of SMAD3 in Alk1 wild type (*Acvrl1*^flox/flox^) MLECs (**Fig. 2B, left panel**). In *Acvrl1*^-/-^ MLECs, BMP9-induced SMAD3 activation was essentially absent, while BMP9-induced activation of SMAD1 was reduced in magnitude and inhibited by LDN-193189 at concentrations ≥ 100 nM, consistent with the low nanomolar potency of LDN-193189 against the remaining BMP type I receptors Alk2, Alk3 and Alk6.^47,48,55^ The potent pharmacologic inhibition of BMP9 signaling by LDN-193189 in BAECs and *Acvrl1*^flox/flox^ MLECs implicate BMP type I receptors for the activation of SMAD1/5 and SMAD2/3; the loss of BMP9-induced SMAD3 activation in *Acvrl1*^-/-^ MLECs demonstrate that ALK1 is essential for SMAD3 activation by BMP9, while the attenuation of SMAD1/5 activation in *Acvrl1*^-/-^ MLECs demonstrate that ALK1 functions in a partly redundant manner with other BMP type I receptors to activate SMAD1/5.

**Figure 2.**
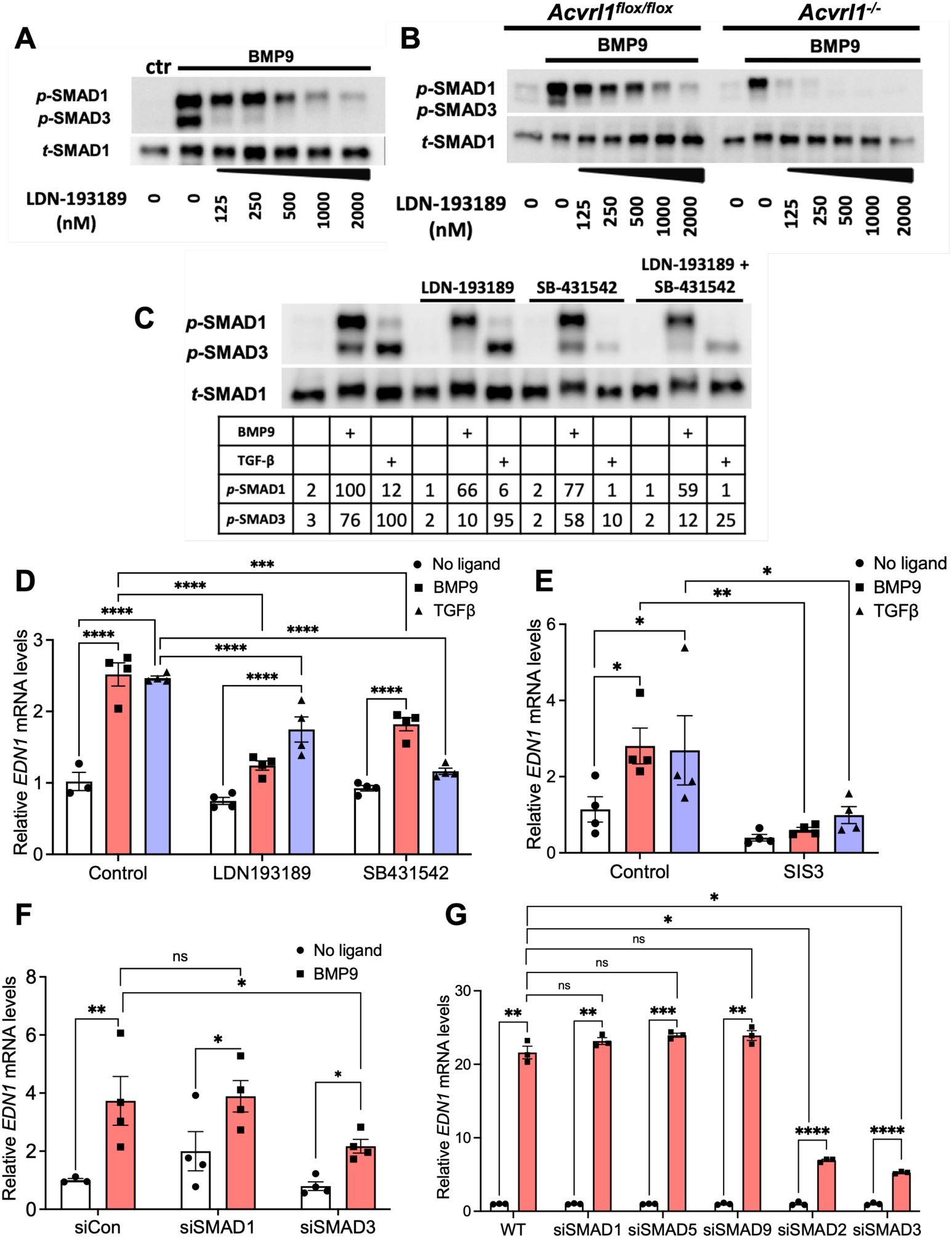
BMP9-mediated ET-1 is dependent upon SMAD2 and SMAD3 activation via ALK1 signaling. (A) Immunoblot of BAECs treated with BMP9 (5 ng/mL) revealed differential sensitivity of SMAD1 vs. SMAD3 activation (*p*-SMAD1, *p*-SMAD3) to ALK1/2/3/6 kinase inhibition with varying concentrations of LDN-193189, vs. total SMAD1 (*t*-SMAD1). (B) Immunoblot of matched ALK1 wild-type and knockout (*Acvrl1*^flox/flox^ and *Acvrl1*^-/-^) mouse microvascular lung ECs (MLECs) treated with BMP9 (5 ng/mL) demonstrate absence of BMP9-mediated SMAD3 activation in ALK1 knockout cells, diminished SMAD1 activation, and more potent inhibition of SMAD1 activation by LDN-193189 in the absence of ALK1. (C) Immunoblot of BAECs stimulated with BMP9 (25 ng/mL ) or TGFβ1 (5 ng/mL) revealed high sensitivity of BMP9-mediated SMAD3 activation and partial sensitivity of SMAD1 activation in response to ALK1/2/3/6 inhibition (250 nM LDN-193189), and modest sensitivity of BMP9-mediated SMAD3 activation and no sensitivity of SMAD1 activation in response to ALK4/5/7 inhibition (5 μM SB-431542), while activation of SMAD3 or SMAD1 by TGFβ1 was sensitive to ALK4/5/7 inhibition. (D) *Edn1* mRNA expression measured by RT-PCR in BAECs stimulated with BMP9 (25 ng/mL) or TGFβ1 (5 ng/mL) revealed differential sensitivity of BMP9- vs. TGFβ1-medated *Edn1* expression to ALK1/2/3/6 (250 nM LDN-193189) or ALK4/57 inhibition (5 µM SB-431542) inhibition. (E) Treatment of BAECs with SMAD3 activation inhibitor SIS3 (5 μM) prevented both BMP9- and TGFβ1-induced expression of *Edn1* mRNA by RT-CR. (F) BMP9-mediated expression of *Edn1* mRNA in BAECs was inhibited by pretreatment with siRNA specific for SMAD3 but not SMAD1 by RT-PCR. (G) BMP9-induced expression of *EDN1* mRNA in TIME cells was inhibited by pre-treatment with siRNA specific for SMAD2 or SMAD3, but not siRNA specific for SMAD1, SMAD5, or SMAD9. Bars represent mean ± SEM, n=3-4 as shown, \**p*<0.05, \*\**p*<0.01, \*\*\**p*<0.001, \*\*\*\**p*<0.0001, 2-Way ANOVA with Holm-Sidak Test (D, G) or Fisher’s LSD test (E, F).

The receptors responsible for BMP9-mediated activation of SMAD1/5 versus SMAD3 were analyzed by immunoblot using small molecule inhibitors of BMP and TGFβ type I receptor signaling in macrovascular and microvascular ECs (**Figs. 2C**, **S2A**). LDN-193189 (250 nM), partially inhibited SMAD1 phosphorylation and essentially abrogated SMAD3 phosphorylation in response to BMP9 in BAECs (**Fig. 2C**) and TIME cells (**Fig. S2A**). Addition of LDN-193189 partially inhibited TGFβ1-mediated activation of SMAD1, but not SMAD3, consistent with the high selectivity of LDN-193189 for BMP type I receptors at this concentration. Treatment of BAECs or TIME cells with SB-431542 (5 µM) modestly impacted BMP9-mediated SMAD1 and SMAD3 activation. Treatment with SB-431542 inhibited TGFβ1-induced activation of SMAD3, and when present, SMAD1 (**Figs. 2C**, **S2A**). Combined treatment with LDN-193189 and SB-431542 did not result in greater suppression of SMAD1 or SMAD3 activation by BMP9 than treatment with LDN-193189 alone, respectively, nor did it result in greater suppression of TGFβ1-induced activation of SMAD3 or SMAD1 by SB-431542 alone. These data in macrovascular and microvascular ECs suggest BMP9-mediated SMAD1 and SMAD3 activation depends primarily on BMP type 1 receptors (ALK1, ALK2, ALK3, or ALK6), while TGFβ/activin type I receptors (ALK4, ALK5, or ALK7) may contribute modestly to BMP9-mediated SMAD3 activation. For TGFβ1, SMAD3 activation requires TGFβ type I receptors, whereas SMAD1 activation has a partial contribution from BMP type I receptors.

Treatment of BAECs with LDN-193189 (125 nM) and to a lesser extent SB-431542 (5 µM) attenuated BMP9-induced *EDN1* expression, while treatment with SB-431542, and to a lesser extent LDN-193189 attenuated TGFβ1-mediated *EDN1* expression (**Fig. 2D**). These data suggested that both BMP and TGFβ type I receptors contribute to the effects of BMP9 and TGFβ1 in promoting *EDN1* expression. The ability of LDN-193189 to block BMP9- and TGFβ1-mediated SMAD1/5 activation and BMP9-mediated SMAD3 activation, and the ability of SB-431542 to block BMP9-mediated SMAD3 activation partially and inhibit TGFβ1 SMAD3 activation (**Fig. 2C**), suggested that cooperation between BMP and TGFβ receptors and their downstream effectors SMAD1/5 and SMAD3 contribute to *EDN1* transcription. While it is known that BMP9 can recruit the activation of BMP-associated SMAD1/5 and TGFβ-associated SMAD3,^14^ their overlapping contributions suggest that SMAD1/5 and SMAD3 effectors may be integrated to regulate *EDN1*.

### BMP9-mediated expression of ET-1 requires SMAD2 and SMAD3 activation

Specific Inhibitor of SMAD3 (SIS3), a small molecule shown to inhibit TGFβ1-induced phosphorylation of SMAD3 and its interaction with SMAD4,^50^ was used to probe the role of SMAD3 in *EDN1* transcription. Treatment of BAECs with SIS3 completely abrogated BMP9-induced as well as TGFβ-induced *EDN1* mRNA expression in BAECs by quantitative RT-PCR (**Fig. 2E**). Similarly, siRNA knockdown of SMAD3 expression in BAECs attenuated BMP9-mediated *EDN1* expression, while siRNA against SMAD1 had minimal effect compared with control siRNA (**Fig. 2F**). In TIME cells, siRNA against SMAD1, SMAD5, or SMAD9 did not impact BMP9-mediated expression of *EDN1*, but siRNA against SMAD2 or SMAD3 potently suppressed this function (**Fig. 2G**). These data suggest SMAD2/SMAD3 are critical nodes for BMP9-mediated regulation of *EDN1* in microvascular and macrovascular ECs.

### BMP9 regulates SMAD3 via ALK1, BMPR2 and ENG in HPAECs

The receptor utilization for BMP9-mediated activation of SMAD1 versus SMAD3 was investigated by immunoblot. Treatment of HPAECs with siALK1 attenuated SMAD1 phosphorylation, and abrogated SMAD3 phosphorylation in response to BMP9 in HPAECs (**Figs. 3A, S3A**), consistent with findings in *Acvrl1*^-/-^ MLECs (**Fig. 2B**), whereas treatment with siALK2 or siALK3 had minimal impact (**Figs. 3A, S3A**). Treatment of HPAECs with siBMPR2 abrogated BMP9-mediated SMAD3 phosphorylation, but did not attenuate SMAD1 phosphorylation (**Figs. 3B**, **S3B**). While previous reports found partially redundant function of endothelial BMPR2 and ACVR2A in transducing BMP9,^14,56^ siACVR2A did not impact BMP9 activation of SMAD1 or SMAD3 (**Figs. 3B, S3B**). Silencing ENG in HPAECs diminished the activation of SMAD3 by BMP9, while silencing ALK5 had minimal effect (**Figs. 3C, S3C**).

**Figure 3.**
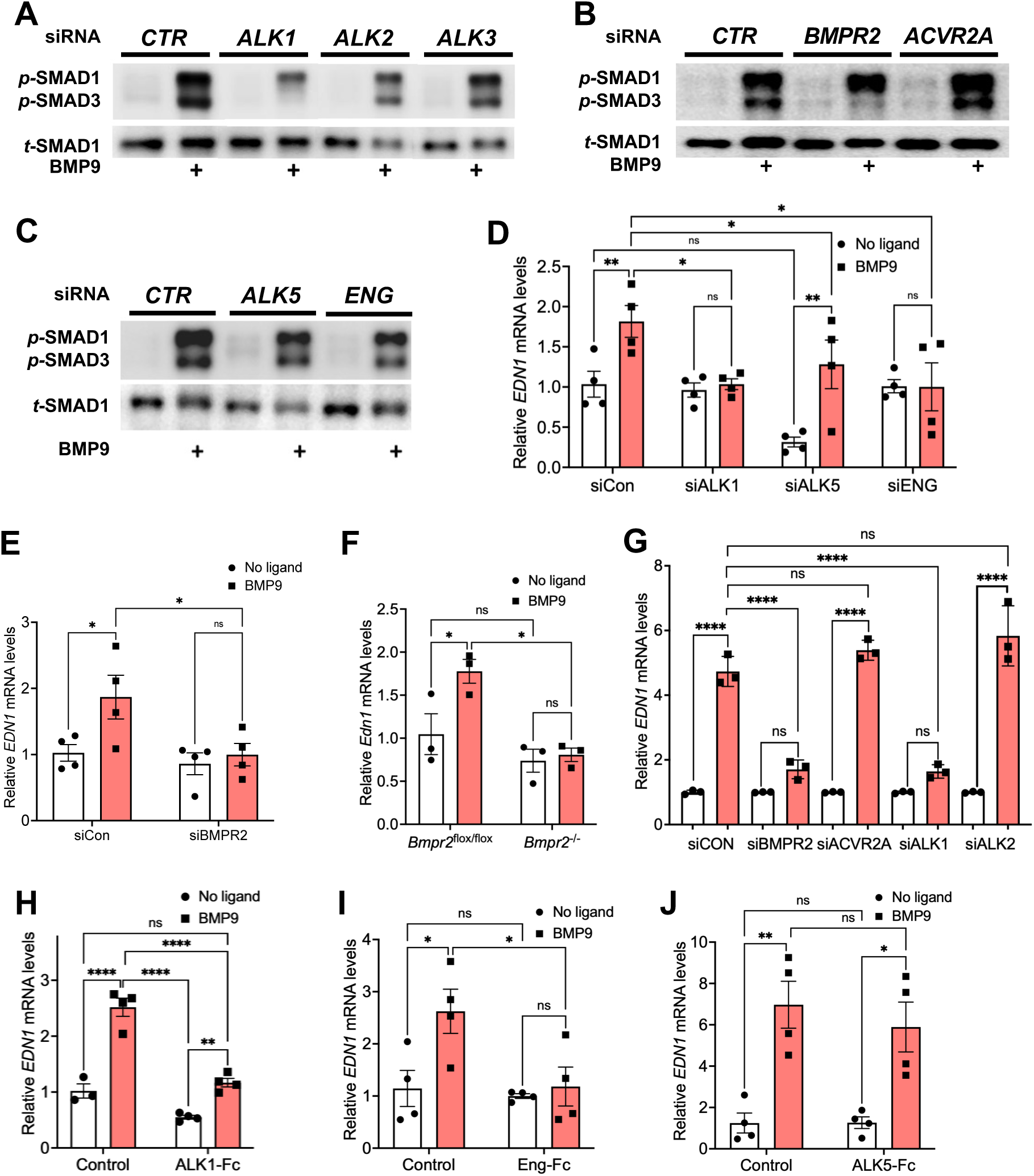
BMP9-mediated *Edn1* expression and SMAD3 activation require ALK1 and BMPR2 signaling. (A-C) Immunoblot of HPAECs following siRNA-mediated knockdown of various BMP type I (*ALK1, ALK2, ALK3*), type II receptors (*BMPR2, ACVR2A*) and co-receptors (*ENG*) revealed that ALK1 and BMPR2 are the primary mediators of SMAD3 activation in response to BMP9 (25 ng/mL), with little contribution from ALK2, ALK3, ALK5, ENG, or ACVR2A, and revealed ALK1 and the primary mediator of SMAD1 activation. (D) RT-PCR of HPAECs pre-treated with specific siRNA revealed inhibition of incremental BMP9-induced *EDN1* expression by si*ALK1* and si*ENG*, but only reduction of basal *EDN1* expression by si*ALK5*. (E) *BMPR2* knockdown in HPAECs abrogated BMP9-mediated *EDN1* expression by RT-PCR. (F) RT-PCR analysis of MLEC revealed loss of BMP9-induced (25 ng/mL) expression of *Edn1* mRNA in *Bmpr2* KO (*Bmpr2*^-/-^) MLEC vs. wild-type MLECs (*Bmpr2*^flox/flox^). (G) RT-PCR analysis of TIME cells pre-treated with specific siRNA revealed loss of BMP9-induced (1 ng/mL) expression of *EDN1* mRNA following knockdown of BMPR2 and ALK1, but not ACVR2a or ALK2. (H-J) Pre-incubation of HPAECs with receptor extracellular domains fusion proteins (E) ALK1-Fc, (F) Eng-Fc, but not (G) ALK5-Fc inhibited BMP9-induced expression of *EDN1* mRNA. Values are mean ± SEM, n=3-4, \**p*<0.05, \*\**p*<0.01, \*\*\*\**p*<0.0001, 2-Way ANOVA with Sidak’s Test.

### BMP9 regulates *EDN1* via ALK1, ENG, and BMPR2 in HPAECs

Receptors involved in BMP9-mediated *EDN1* expression were interrogated in HPAECs using siRNA for BMP/TGFβ receptors and co-receptors ALK1, ALK5, ENG, and BMPR2. Expression of genes were reduced by 61%-95% by their respective siRNAs, confirming efficient silencing (**Fig. S3D**). Compared to control siRNA, siALK1 and siENG abrogated BMP9-induced *EDN1*, whereas siALK5 reduced basal *EDN1* but did not disrupt BMP9-induced *EDN1* (**Fig. 3D**). Treatment with siBMPR2 abrogated BMP9-induced *EDN1* (**Fig. 3E**). Similarly, BMP9-induced Edn1 was disrupted in conditional (*Bmpr2*^-/-^) but not WT (*Bmpr2^flox/flox^*) MLECs (**Fig. 3F**). Using TIME cells to confirm these findings in human microvascular ECs, siBMPR2 and siALK1 both attenuated BMP9-induced *EDN1*, whereas siACVR2A and siALK2 had no impact **(Fig. 3G**). Taken together, these data show that BMPR2 and ALK1 are essential for BMP9-mediated SMAD3 activation and *EDN1* expression, which was found to require SMAD2 and SMAD3 activation. ENG also contributed to BMP9-induced SMAD3 activation and was essential for BMP9-induced *EDN1*, while ALK5 contributed to basal but not BMP9-induced *EDN1* expression. Using pharmacologic inhibitors, siRNA for receptors and SMAD effectors, these experiments demonstrate BMP9 regulation of *EDN1* in microvascular and macrovascular ECs requires the function of BMPR2, ALK1, and ENG to activate SMAD3, with potential participation of ALK5.

### ALK1 and ENG, but not ALK5 extracellular domains inhibit BMP9-induced *EDN1* expression

BMP9 is known to utilize signaling complexes of ENG and ALK1,^15,16,18^ which is reported to interact with ALK5 in regulating endothelial responses to TGFβ1.^53,54,57^ Partial inhibition of BMP9-induced *EDN1* expression by ALK4/5/7 inhibitor SB-431542 (**Fig. 2D**) prompted testing whether the extracellular domains of these receptors might interact with BMP9. Receptor extracellular domains expressed as IgG Fc-fusion proteins ALK1-Fc and ENG-Fc, but not ALK5-Fc attenuated BMP9-induced *EDN1* expression in BAECs (**Fig. 3H-J**), consistent with the higher affinity of BMP9 for ALK1 and Eng as compared with ALK5 and other BMP/TGFβ type I and type III receptors.^52^

### BMP9 requires a putative nuclear factor-1 (NF-1)-binding element for regulating ET-1

Previous reports show that upstream of the human *ET-1* transcription initiation site, an AP-1 site (-102/-108 bp) and TGFβ-responsive SMAD2/3-SMAD4 binding elements (SBE1 and SBE2, -171/-193) act cooperatively to regulate TGFβ1-induced transcription of *EDN1*.^58^ We tested whether these or additional elements might be required for BMP9-mediated regulation of *EDN1*. Using reporter constructs containing truncated 5’ promoter segments of varying length, we found a minimum promoter of -298 bp from the transcription initiation site was required for BMP9-stimulated transcription of ET-1, whereas a -277 bp upstream segment was sufficient for TGFβ1 but not BMP9 activity in BAECs (**Fig. 4A**). Analysis of the region between -277 and -298 bp revealed the presence of a putative NF-1 binding site at -293 bp, which enhanced but was not essential for TGFβ1-mediated *EDN1* transcription.^34^ This putative NF-1 domain bears homology to a BMP9-sensitive SMAD1/5 response element called “MEME2” previously identified by CHIP-Seq in BMP9-stimulated human umbilical vein endothelial cells (**Fig. 4B**).^59^

**Figure 4.**
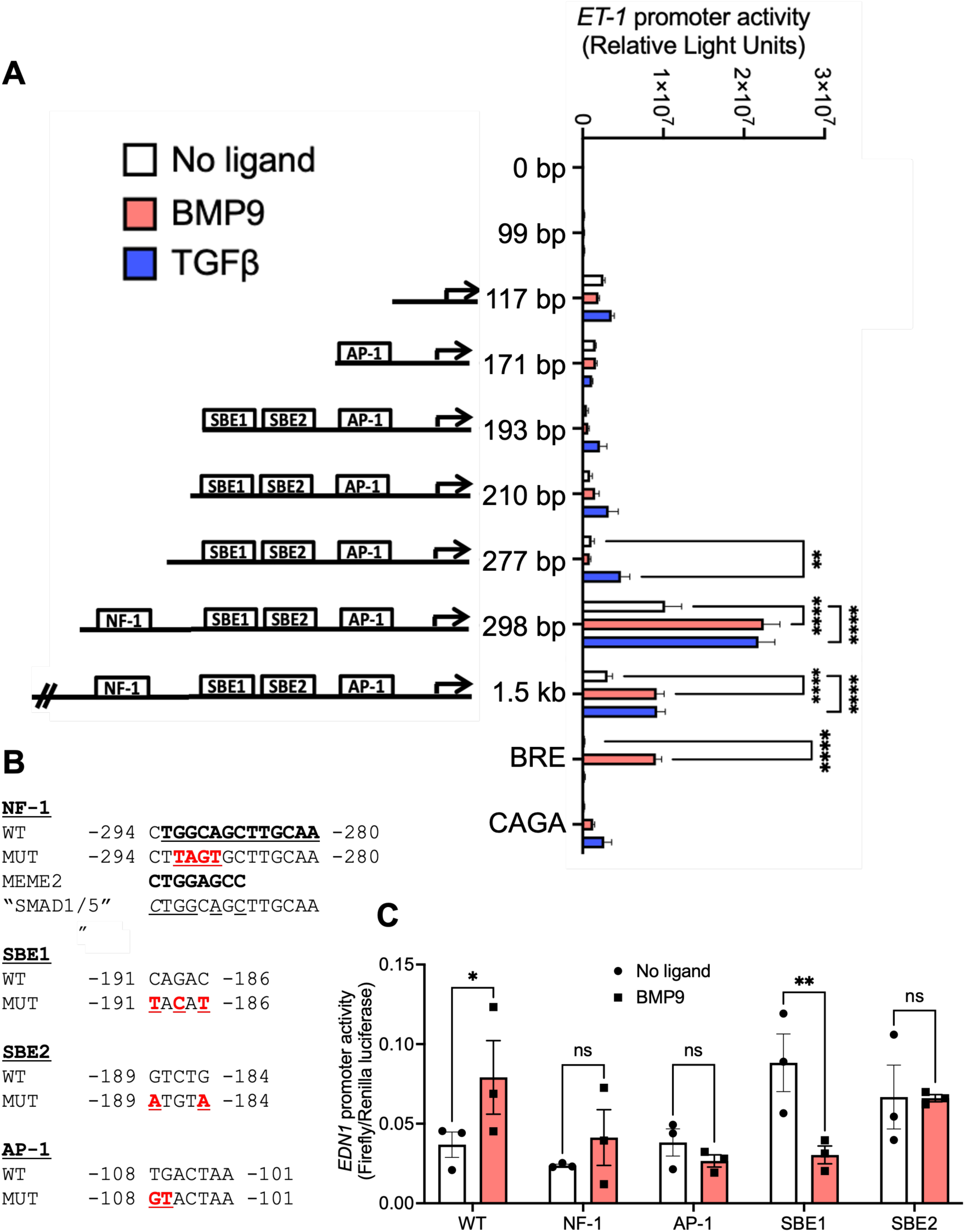
BMP9-induced *EDN1* promoter activity requires the presence of a putative Nuclear Factor-1 (NF-1) binding element with sequence identity to a SMAD1/5 binding site. *EDN1* promoter activity induced by BMP9 versus TGFβ1 in BAECs was assayed using either wild type human *EDN1* promoter of varying lengths or using a full-length promoter with mutations in BMP- and TGFβ-specific binding elements. (A) Previously described SMAD2/SMAD3 binding element (SBE1/SBE2), activator protein-1 (AP-1) binding site, and a putative NF-1 site with homology to a consensus SMAD1/5 binding element (“MEME2”) were required for *EDN1* transcription in response to BMP9, whereas TGFβ only required the SBE1/SBE2 and AP-1 sites. (B-C) Mutant constructs were generated within a -650 bp version of the *EDN1* promoter reporter construct and compared with the signaling of the wild-type construct by luciferase assay. Cells were deprived of fetal calf serum (FCS) for 16 h, and then treated with 25 ng/mL of BMP9 or 5 ng/mL of TGFβ1 for 12 h. Values are mean ± SEM, n=3-5, \**p*<0.05, \*\**p*<0.01, \*\*\**p*<0.001, \*\*\*\**p*<0.0001, 2-Way ANOVA with Sidak’s Test, or in the case of promoter mutants, by Fisher’s exact test for each mutant / wild-type pair.

To confirm the role of the putative NF-1 binding site in BMP9-mediated activity as well as previously identified sites known to facilitate TGFβ1-mediated activity, mutant promoter constructs based on the -650 bp wild-type *EDN1* promoter were used. Point mutations of critical residues in the SBE1, SBE2, AP-1, as well as putative NF-1 binding sites all attenuated the increased activity of BMP9 (**Fig. 4B-C**), demonstrating BMP9 required the putative NF-1 site in addition to AP-1 and SBE1/2 sites previously identified as essential for TGFβ1. These observations are consistent with the requirement of SMAD2/3 activation and synergistic effect of SMAD1/5 activation in BMP9-induced *EDN1* expression, and is compatible with the disruption of BMP9-mediated *EDN1* expression with LDN-193189 tretment, or ALK1 knockout or knockdown, all of which disrupt SMAD1/5/9 and SMAD2/3 activation (**Fig. 2B-D**). Taken together, these observations may be explained by the participation of NF-1 as a SMAD1/5 binding element enhancing both BMP9- and TGFβ-mediated *EDN1* transcription (**Fig. 4A**).

## DISCUSSION

Given the broad importance of BMP9 in vascular patterning and endothelial function,^23,25,27,28,60–66^ and of endothelial ET-1 in regulating diverse aspects of vascular tone, homeostasis, angiogenesis, fibrosis and matrix remodeling, the BMP9-ET-1 signaling axis holds relevance as a critical effector mechanism bridging BMP9 to vascular tone and remodeling.^36,67–73^ Our current study sought to pinpoint molecular mechanisms by which BMP9 regulates ET-1 using complementary small molecule inhibitor, ligand trap, receptor- and SMAD- specific siRNA, conditional *Alk1* and *Bmpr2* knockout, and *EDN1* promoter analysis strategies. The study employed primary human PAECs, murine PMVEC, BAEC, and human TIME cells to model diverse macrovascular and microvascular beds of the pulmonary and systemic circulation, leveraging genetic tractability of murine EC and efficient gene transfer in bovine and immortalized human EC. Consistent findings were obtained despite the diverse anatomic origins, species, and primary vs. transformed status of these EC, suggesting the mechanisms by which BMP9 regulates ET1 and this function itself are highly conserved.

BMP9 is one of several ligands of the TGFβ family that activates both SMAD1/5/9 and SMAD2/3 signaling proteins, two sets of effectors that in various contexts exert opposing biological consequences on cell differentiation, hypertrophy, and extracellular matrix remodeling.^74^ We found that both SMAD1/59 and SMAD2/3 signaling required the same ternary receptor complex of BMPR2, ALK1, and ENG, all highly enriched in endothelium. While previous studies supported the role of ALK1 in BMP9-mediated transcriptional responses,^14,29^ and other studies suggested that ALK2 drives BMP9-induced *EDN1* in the absence of BMPR2,^37^ our study found that ALK1 is essential for *EDN1* transcription. While other studies reported that ALK5 and ENG were not involved in mediating BMP9 responses,^14^ here we found ENG to be indispensable. In contrast to a previous study which attributed BMP9 activation of SMAD1/5/9 to BMPR2, and SMAD2 to ACVR2A, and no activation of SMAD3,^14^ we found that BMPR2-ALK1-ENG signaling was essential for BMP9-mediated activation of SMAD2 and SMAD3, and a major driver of SMAD1/5/9, challenging the concept that ALK1 only recruits SMAD1/5/9. Park and colleagues used an siRNA approach to show that loss of endothelial *BMPR2* attenuated BMP9-mediated *EDN1* expression, but with modest increases in basal pre-pro-*EDN1* mRNA and peptide expression.^29^ In that study, BMP9-induced *EDN1* expression required activity of BMPR2 and ALK1, as observed in the present study, but in contrast to our findings, the effects were attributed to SMAD1 and MAPK p38 activation rather than SMAD3, the role of which was supported in this study by multiple approaches. The importance of SMAD3 in regulating tissue fibrosis and extracellular matrix production has been described in the context of TGFβ/ALK5 signaling,^45,53^ and as a mechanism of TGFβ-mediated regulation of ET-1, whereas linking the BMP9-BMPR2-ALK1-ENG signaling complex to ET-1 via SMAD3 represents to our knowledge the first physiologic function of BMP9 ascribed to its non-classical activation of SMAD3. Our current results attach a novel and important function to “lateral activation” of SMAD3 by ALK1, and a mechanism for the pleiotropic functions of BMP9. Whether the combined activation of SMAD1/5 and SMAD3 by BMP9 regulates similar or distinct fibrosis and matrix remodeling programs as TGFβ is a subject for further study.

Consistent with previous reports, we found that BMP9 elicited ET-1 expression with similar potency to TGFβ1 in microvascular and macrovascular EC linages. While TGFβ1 is known to promote *EDN1* transcription via cooperation of AP-1 and SBE promoter domains downstream of ALK5 and SMAD3,^33–35,58^ we found BMP9 requires an additional putative NF-1 or SMAD1/5 binding domain, the presence of which enhances TGFβ and BMP9 activity (**Fig. 5**). We linked phosphorylation of SMAD3 to ALK1 function using conditional ALK1-deficient ECs, siRNA specific for ALK1, and LDN-193189, a selective antagonist of ALK1/2/3/6 that was used at concentrations sufficient to inhibit ALK1 without inhibiting ALK4/5/7.^47,75^ Potential integration of SMAD1/5/9 and SMAD2/3 activation by *EDN1* promoter elements lends further evidence that pleiotropic effects of BMP9 may hinge on activating both BMP and TGFβ effectors.

**Figure 5.**
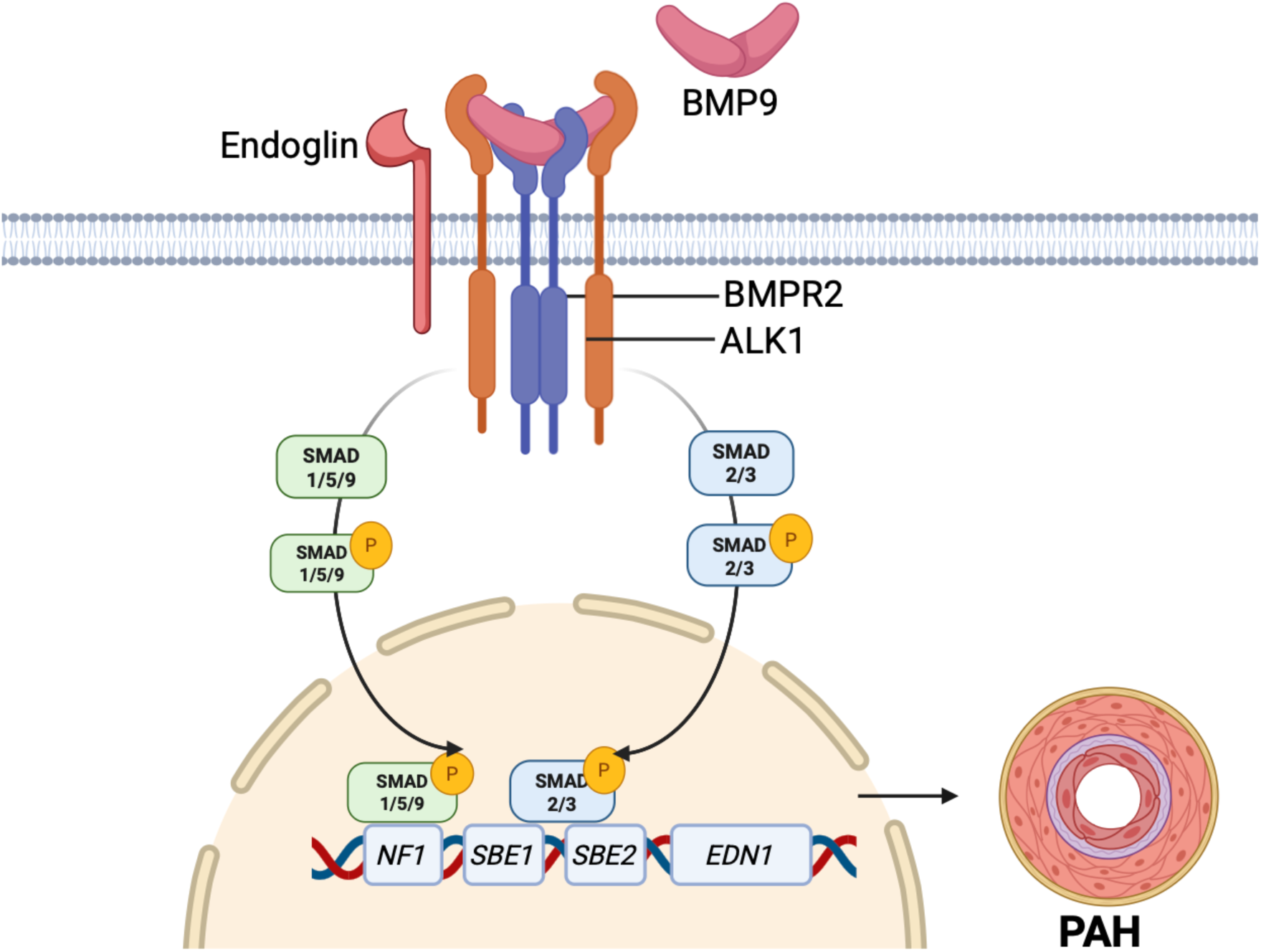
Graphical abstract. The schematic illustrates the proposed mechanism demonstrated in this study. BMP9 initiates the formation of a membrane receptor complex consisting of two type I receptors (ALK1) with two type II receptors (BMPR2) and two ENG coreceptors, promoting transphosphorylation of the ALK1 kinase. Activated ALK1 then phosphorylates SMAD 2/3 and SMAD 1/5/9. Subsequently, pSMAD 2/3 and pSMAD 1/5/9 translocate to the nucleus where they regulate *EDN1* transcription via binding to SBE1/SBE2 and NF1 promoter sites, respectively, in microvascular and macrovascular endothelial cells, thereby promoting PAH.

The traditional model of BMP/TGFβ receptor activation does not readily explain how BMP type I receptors may activate SMAD2 or SMAD3, typically considered substrates for TGFβ and activin receptors ALK4/5/7. A converse type of “lateral activation” of SMAD1/5/9 by TGFβ has been attributed to either i) dual specificity of TGFβ type I receptor ALK5 for SMAD1/5/9 and SMAD2/3, ii) mixed BMP/TGFβ receptor complexes; or iii) mixed SMAD complexes as potential mechanisms.^53,76,77^ In the current study TGFβ type I and type II receptors played lesser roles than ALK1 in the activation of SMAD3 by BMP9, and affected basal levels of ET-1 expression rather than incremental expression due to BMP9. Combining SB-431542 with LDN-193189 did not inhibit BMP9-induced SMAD3 activation beyond LDN-193189 alone, whereas ablation of ALK1 abrogated BMP9-induced SMAD3 activation and ET-1 expression, making contributions of other BMP/TGFβ type I receptors non-essential. Here, ALK1 appeared to play a direct role in the activation of SMAD3 and ET-1 regulation, suggesting dual specificity of ALK1 for SMAD2/3 in addition to BMP-associated SMADs. The requirement for SMAD2/3 and putative SMAD1/5 binding elements for BMP9 activity (**Fig. 5)**, and the inhibition of BMP9-mediated SMAD1 and SMAD3 activation, and *EDN1* expression by LDN193189, suggest a common mechanism of SMAD1/5 and SMAD2/3 activation, and their cooperation in *EDN1* transcription (**Fig. 2C-D**), While treatment with siSMAD1, siSMAD5, or siSMAD9 individually did not inhibit BMP9-mediated *EDN1* transcription (**Fig. 2F-G**), redundant function of these SMADs cannot be ruled out. Combinations of siSMAD1/5 and siSMAD1/5/9 were also tested, but led to loss of EC viability, and thus necessity of SMAD1/5/9 as a group cannot be ruled out by this approach.

Explanations for why SMAD2/3-binding (SBE1/2) and AP-1 elements in the *EDN1* promoter sufficient for TGFβ1 were insufficient for BMP9 could lie in differential amplitude, duration, or subcellular localization of SMAD3 activation due to BMP9 vs. TGFβ1 signaling, requiring the putative NF-1/ SMAD1/5 site in addition. An identical binding motif was identified by Morikawa et al. in a genome-wide study of SMAD1/5-targeted promoters within BMP9-responsive genes in human umbilical vein endothelial cells (HUVECs) using chromatin immunoprecipitation followed by sequence analysis (ChIP-Seq).^42^ Of five sequence motifs identified in this study, the MEME2 motif (CTGGAGCC) shows 6/8 (75%) identity with the putative NF-1 binding site in the human *EDN1* promoter (**Fig. 4B**). In addition, analysis of SMAD1/5-binding sites in the associated NCBI Sequence Read Archive (accession number SRA030442) revealed 12 reads of SMAD1/5-binding sites containing the putative NF-1 element of the *ET-1* promoter, (TGGCAGCTTGCAA), as well as 7 reads containing an additional C at the 5′ terminus, both of which are identical in sequence to the BMP9-responsive segment of the *ET-1* promoter at -294 bp. Thus, the high homology or identity of the putative NF-1 site with several ChIP-Seq-identified BMP9-responsive SMAD1/5 binding sites, and its close proximity to the SMAD3-SMAD4-binding sequence (SBE1/SBE2) provide a potential mechanism by which SMAD1/5 and SMAD3 signaling might cooperate. This cooperativity may represent a motif for the regulation of other vasoactive endothelial gene products requiring engagement of both BMP and TGFβ effectors and is a subject for future detailed studies.

It should be noted that ET-1, like BMP9, may have opposing, context-sensitive functions. While ET_A_ receptor signaling by ET-1 in smooth muscle promotes vasoconstriction and fibrosis, ET_B_ receptor negative feedback signaling in endothelium promotes vasodilation, natriuresis, anti-proliferative and anti-inflammatory effects, in this manner regulating endothelial quiescence similar to what has been proposed for BMP9.^18,71^ Autocrine and paracrine effects of BMP9-ET-1 signaling may thus impact adjacent vascular media and endothelium with consequences that depend on the vascular bed, tissue, and local hemodynamic and inflammatory state.

While the current findings potentially link BMP9 to regulation of vascular tone via ET-1/ET_A_ signaling, as highlighted by protection of *Gdf2^-/-^* mice from experimental PH, and the inhibition of BMP9-mediated vasoconstriction in chorioallantoic membranes by ERA,^38,39^ these findings may also explain PH-associated vascular remodeling. Our recent single nucleus RNA-seq analysis of experimental PH revealed that antagonism of BMP9 inhibits the PH-associated reprogramming of gCap EC to arterial EC,^40^ a cellular transition that has been observed in genetic models of PH and human PAH.^78^ Expansion of arterial EC, the EC subset that expresses ET-1 most abundantly in human PAH and experimental PH,^40,51^ may drive muscularization of the distal pulmonary circulation. Thus, beyond regulation of vascular tone, the BMP9-ET-1 axis may provide a mechanistic link between dysregulated BMP signaling and vascular remodeling in PAH.

The efficacy of inhibiting BMP9 in experimental PH, the potential role of BMP9 as a driver of vascular tone and EC phenotype modulation, all likely to involve regulation of endothelial production of ET-1, have served as the rationale for a Phase 2 proof-of-concept trial testing an anti-BMP9 monoclonal antibody in clinical PAH (ClinicalTrials.gov NCT06137742).^41^ Given the broad impact of ET-1 in vascular homeostasis and remodeling, and the equally broad roles BMP9 in EC and vascular function, the novel BMP9-ALK1-SMAD3 signaling motif defined here should be considered in the pathophysiology of other vascular syndromes involving dysregulated BMP/TGFβ signaling.

## Funding

This work was supported by support from the Actelion Entelligence Young Investigator Award (J.B.), the US National Institutes of Health (HL159443, HL131910, HL079943, P.B.Y.; HL007604 J.B. and I.N.; and HL007208, J.B.), the Pulmonary Hypertension Association (P.B.Y), the Leducq Foundation Transatlantic Network of Excellence (P.B.Y.), the Spanish Ministry of Science, Innovation and Universities (PID2022-136703OB-I00) (F.R.P), and the Howard Hughes Medical Institute (P.B.Y.).

## Author Contribution Statement

JB, SS, LT, SPF, IN, YZ, GRG, FRP, and PBY contributed to overall experimental design, contributed data, and participated in the analysis of the data. JB, SS, FRP and PBY drafted the manuscript, while all authors participated in the revision of the manuscript.

## Acknowledgments

The authors wish to thank Drs. K. Bloch, E. Li, P. ten Dijke, and T. Michel for helpful discussions.

## Conflicts of Interest

PBY is a co-founder, consultant, and stockholder for Keros Therapeutics, which develops therapies for cardiovascular, hematologic, and musculoskeletal diseases targeting bone morphogenetic protein and TGFβ signaling pathways. PBY is a co-founder and board member of Modal Therapeutics, which develops therapies for vascular and metabolic disease. PBY also receives research funding from Pfizer, Inc., with which patent applications for PAH-specific therapies have been filed. The interests of PBY are reviewed and managed by Mass General Brigham in accordance with their conflict-of-interest policies.

## Supplementary Data

**Supplemental Figure 1.**
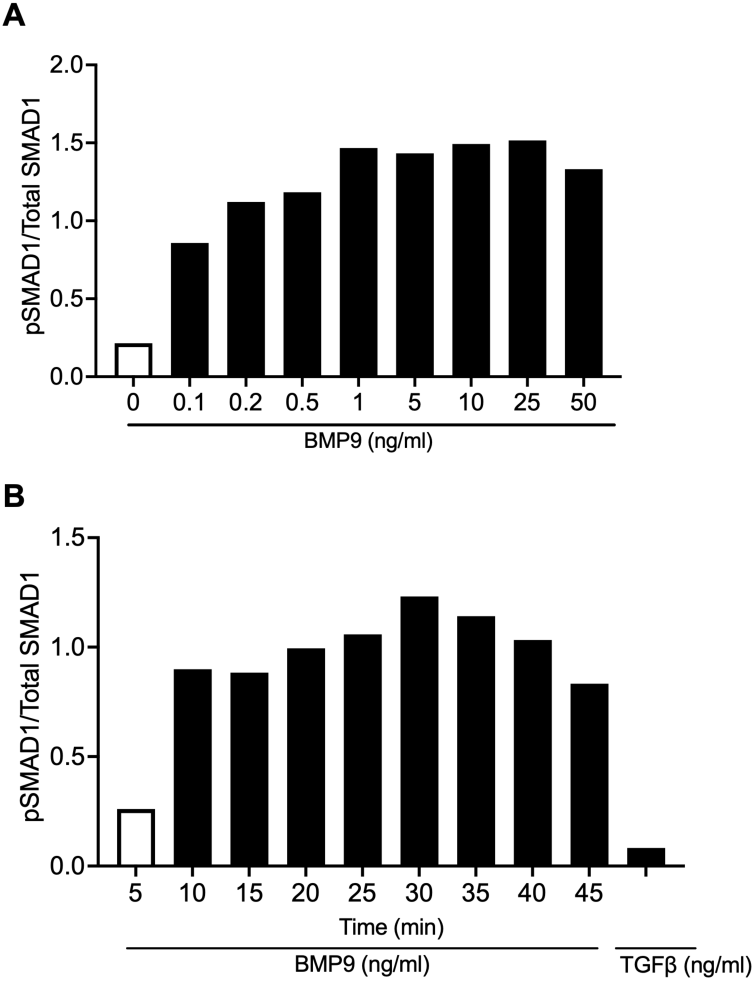
(A). Densitometric graph showing activation of SMAD1 in BAECs stimulated with varying concentrations of BMP9 for 25 min in a dose-dependent manner. (B). Densitometric graph in BAECs stimulated with BMP9 or TGFβ1 shows activation of SMAD1 in a time-dependent manner.

**Supplemental Figure 2.**
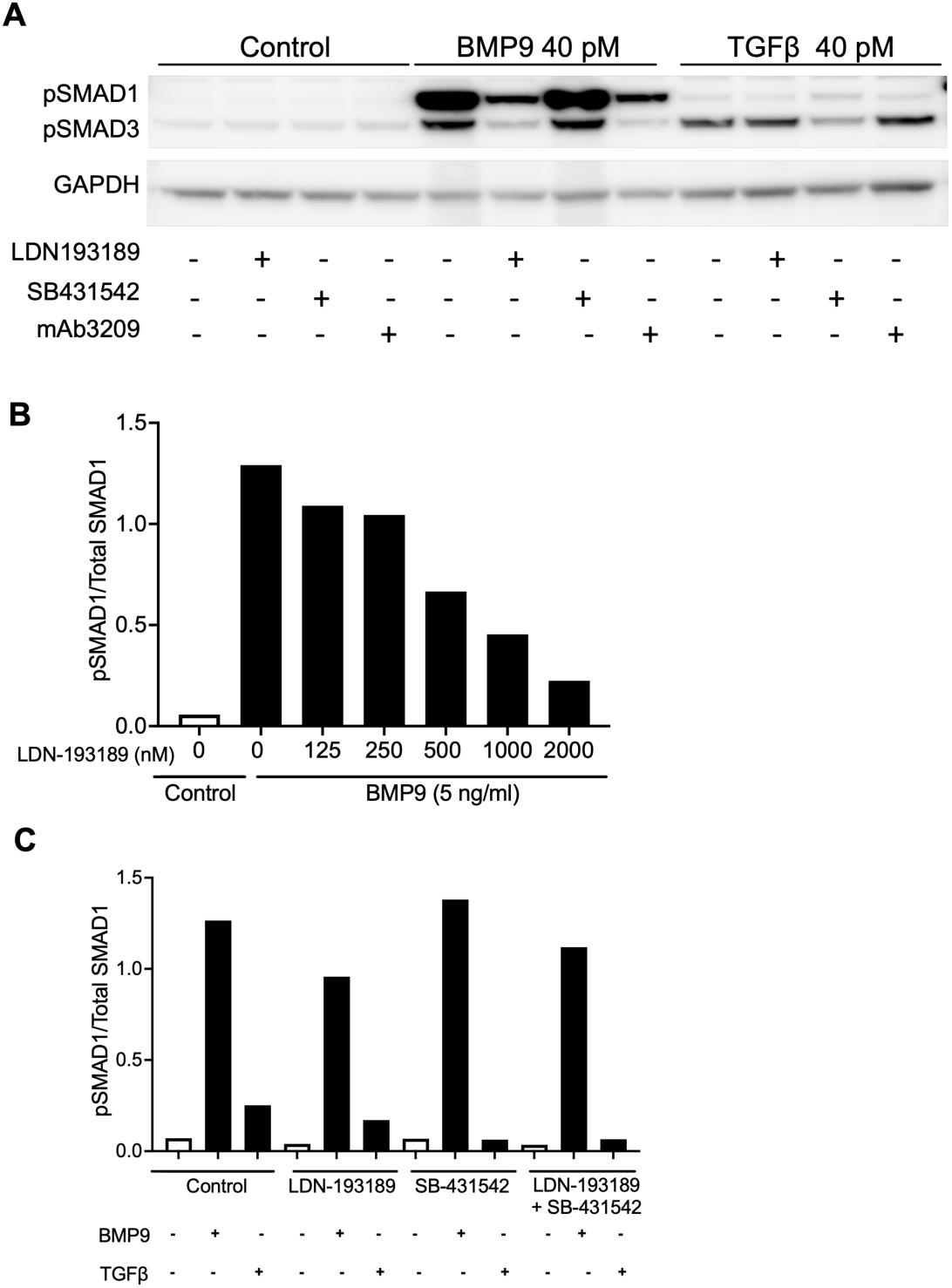
(A) Immunoblots of protein lysates of telomerase-immortalized human microvascular endothelial (TIME) cells pre-incubated with 250 nM LDN-193189 or 5 μM SB-431542 or 40 nM mAB3209, and then stimulated with BMP9 (40 pM) or TGFβ1 (40 pM) showing the expression of SMAD1 and SMAD3 activation compared to untreated controls. (B) Densitometric graph showing activation of SMAD1 in BAECs, pre-incubated with various concentrations of LDN-193189 and then treated with BMP9 (5 ng/mL). (C) Densitometric graph showing the ratio of pSMAD1/total SMAD1 in BAECs pre-incubated with 250 nM LDN-193189 or 5 μM SB-431542, or both, and then stimulated with of BMP9 (25 ng/mL) or TGFβ1 (5 ng/mL).

**Supplemental Figure 3.**
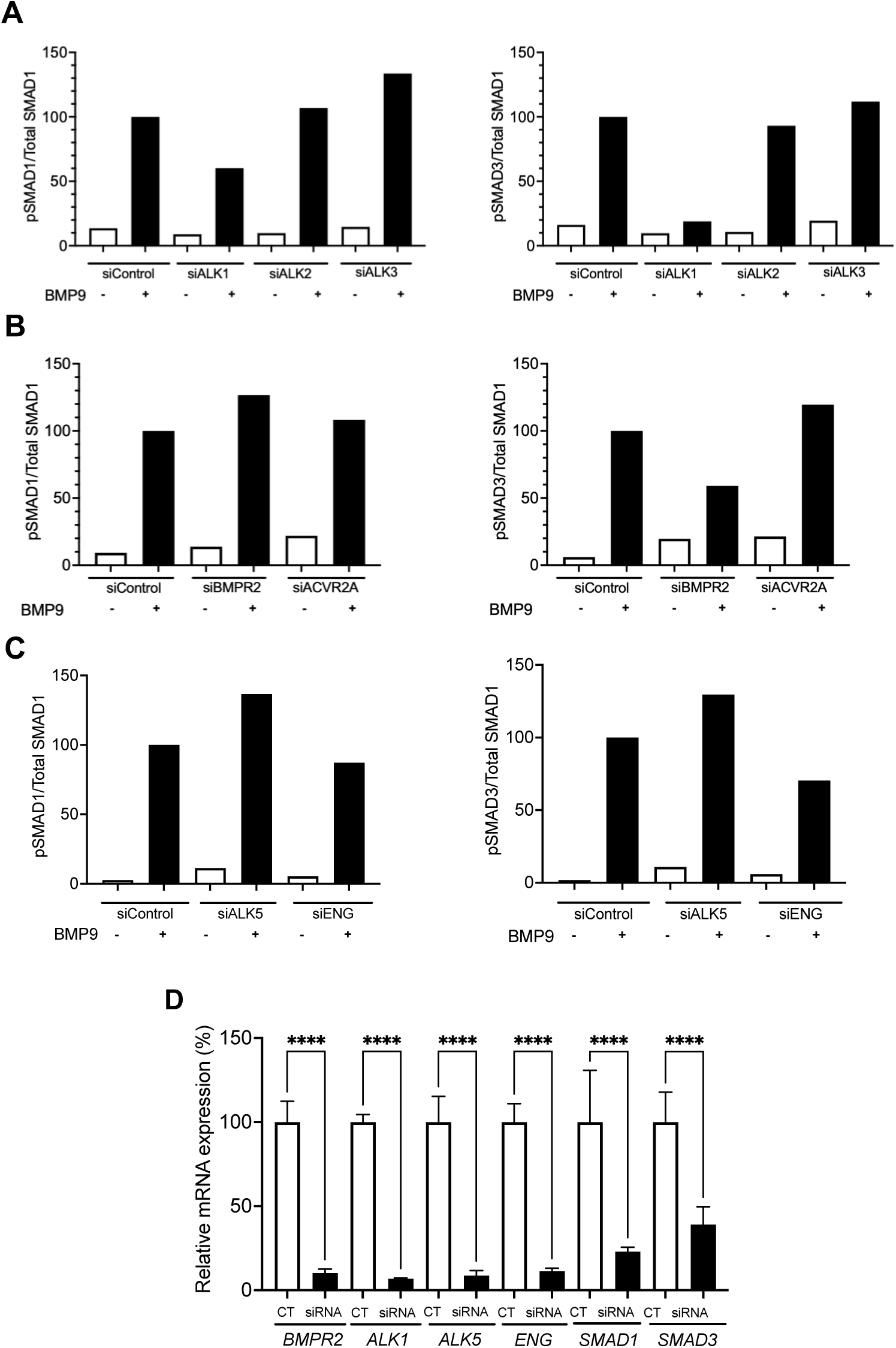
(A-C) Densitometric analyses of immunoblots of HPAECs (see **Fig. 3A-C)** following siRNA-mediated knockdown of various BMP type I (*ALK1, ALK2, ALK3*), type II receptors (*BMPR2, ACVR2A*) and co-receptors (*ENG*) revealed that ALK1 and BMPR2 are the primary mediators of SMAD3 activation in response to BMP9 (25 ng/mL), with little contribution from ALK2, ALK3, ALK5, ENG, or ACVR2A, and revealed ALK1 and the primary mediator of SMAD1 activation. (D) siRNA transfection of HPAEC with siRNAs specific for various receptors, coreceptors, and SMAD effectors (BMPR2, ALK1, ALK5, ENG, SMAD1, SMAD3) confirms 61%-95% knockdown compared to control siRNA. Values are mean ± SD n=5, \*\*\*\**p*<0.0001 by One-Way ANOVA, with Sidak’s test for multiple comparisons.

**Supplemental Table 1.**
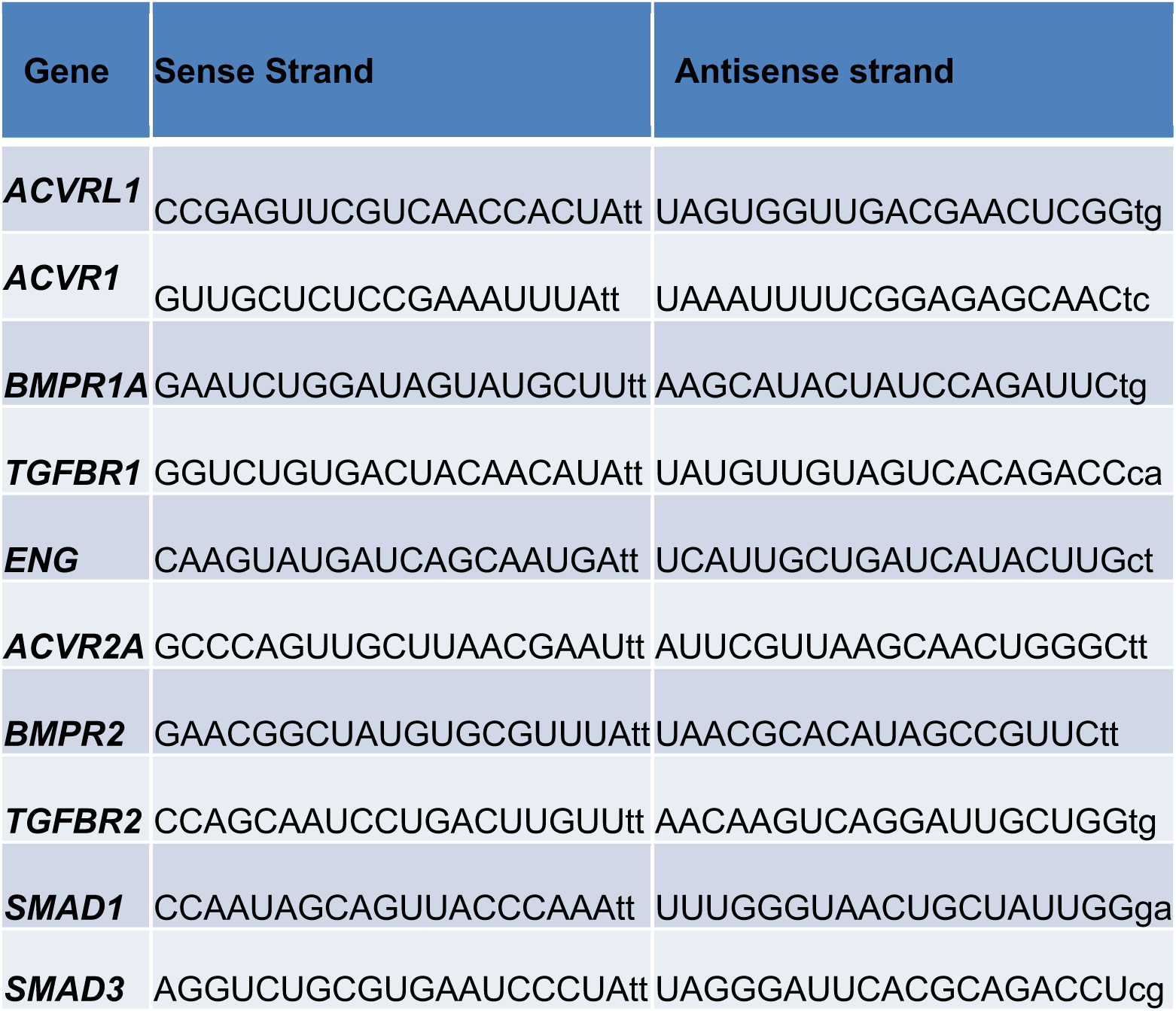
List of small interfering RNA oligonucleotides used in our study to down-regulate expression levels of various members of BMP and TGFβ signaling pathways.

